# Structure-Function Coupling Aligns with a Unimodal-to-Transmodal Gradient in the Mouse Cortex

**DOI:** 10.64898/2025.12.04.692468

**Authors:** Ruixiang Li, Mika Takahashi, Mizuki Kurosawa, Teppei Matsui, Aya Ito-Ishida

**Author notes:** Corresponding authors: Aya Ito-Ishida, M.D., Ph.D.,Laboratory for Brain Development and Disorders, RIKEN Center for Brain Science, 2-1 Hirosawa, Wako, Saitama 351-0198, Japan., Teppei Matsui, Ph.D., Graduate School of Brain Science, Doshisha University, Kyoto, 610-0394, Japan.

## Abstract

Understanding how anatomical connectivity shapes brain activity is essential for clarifying brain function and disorders. In the human brain, the regional heterogeneity in structure-function (S-F) coupling is well characterized by measuring correlations between structural and functional connectivity. However, it remains unclear whether the same principles apply to the mouse cortex, where biological mechanisms can be studied directly. Here, we mapped S-F coupling across the mouse cortex by combining high-resolution structural connectivity derived from axonal tracing with resting-state functional connectivity measured by wide-field calcium imaging. Our findings revealed that structural connectivity imposes a robust yet regionally variable constraint on functional connectivity. As in humans, the S-F coupling was strong in primary sensorimotor areas and weaker in association areas, demonstrating a gradient from unimodal-to-transmodal cortical organization. This spatial variation covaried with intrinsic cellular properties, including myelination, excitation-inhibition balance, synaptic density, and gene expression profiles, but did not align with the anatomically defined cortical hierarchy. Our findings highlight graded S-F coupling as a common organizational principle in both the mouse and human cortex, providing a framework for future mechanistic studies using mouse models.

## Introduction

Behavior arises from the coordinated activity of neuronal ensembles that coactivate across multiple brain areas^1^. Functional connectivity (FC), which measures statistical dependence between neural activity in different regions, has been widely used to characterize these ensembles in health and disease^2–5^. Traditionally, FC in the resting state was thought to reflect structural connectivity (SC), which refers to the strength of anatomical wiring between regions^3,6^. Consistent with this ‘structure-function (S-F) coupling’ hypothesis, studies based on MRI have reported a robust association between SC and FC^7–11^. However, SC and FC are not equivalent: significant mismatches exist, and even sophisticated computational models have shown that SC alone is insufficient to predict FC^12,13^. These discrepancies indicate that additional mechanisms, such as local microcircuitry, polysynaptic interregional connections, and neuromodulatory pathways, are involved in shaping FC beyond direct anatomical connections^14,15^.

Research in humans revealed a pronounced regional heterogeneity in S-F coupling^16,17^. Notably, the degree of S-F coupling varies along a functional gradient that aligns with the brain’s functional organization^18,19^. S-F coupling is strongest in primary visual and motor cortices (unimodal areas) and progressively weakens in prefrontal and parietal cortices (transmodal or multimodal areas). Although the physiological significance of this regional heterogeneity in S-F coupling is not fully understood, the reduced S-F coupling in transmodal areas suggests that these areas may possess distinct biological characteristics essential for flexible and higher-order cognition^20,21^. In support of this view, local S-F coupling in humans has been linked to several intrinsic cortical properties, such as myelination and excitation-inhibition (E/I) balance^22^.

To further clarify the biological mechanisms regulating S-F coupling, it is essential to connect macroscopic connectivity with microscopic cellular properties. However, current MRI-based analyses of S-F coupling face methodological limitations: SC derived from diffusion MRI tractography often overestimates long, smooth anatomical tracts and fails to precisely estimate axonal density^23^. In addition, FC is obtained by functional MRI, which measures slow hemodynamic responses instead of neuronal activity *per se*^24^. These limitations highlight the need for mechanistic studies in mice, where cellular-resolution measurements and causal manipulations are feasible. However, despite recent investigations^25–28^, the regional heterogeneity of S-F coupling remains poorly characterized in the mouse cortex. A key question is whether the S-F coupling in the mouse cortex aligns with the biological gradients that underlie cortical organization^29–32^. Specifically, is the unimodal-to-transmodal decrease in S-F coupling, well described in humans with expansive association cortices, also present in the more compact mouse cortex? Addressing this question is essential for understanding the cellular mechanisms of cortical organization.

In this study, we characterized S-F coupling in the mouse cortex by integrating neurophysiologically grounded measurements of SC and FC. To obtain SC, we used axonal tracing datasets in the Allen Mouse Brain Connectivity Atlas (AMBCA)^29,33,34^. For FC, we conducted wide-field calcium imaging in awake mice. By analyzing these datasets at fine spatial resolutions, we mapped S-F coupling across the mouse cortex. Although SC and FC were strongly correlated, SC explained only a portion of FC, and local S-F coupling varied systematically across cortical areas. These regional differences in S-F coupling correlated with intrinsic cellular properties, including myelination, E/I balance, synaptic density, and gene-expression profiles, but did not track anatomically defined cortical hierarchy. Notably, the spatial pattern of S-F coupling in mice aligned with a functionally defined hierarchical gradient extending from unimodal to transmodal cortex, which is consistent with previous observations in humans. Together, our findings suggest that the region-dependent S-F coupling is a conserved principle of cortical organization in both mice and humans, providing a framework for exploring the cellular and molecular mechanisms that regulate the S-F relationship.

## Results

### Imperfect Correspondence between Structural and Functional Connectivity

We first established high-resolution maps of SC and FC in the mouse cortex. We used a voxel-level SC based on AMBCA dataset^34^, which gives axonal projection density (PD) between regions at 100-µm resolution. **Fig. 1a** shows representative examples of PD originating from a single seed voxel. To align the structural data with the perspective of wide-field calcium imaging, the volumetric cortical SC data were projected as topview maps (**Fig. 1a** & **Supplementary Fig. 1;** see Methods).

**Figure 1.**
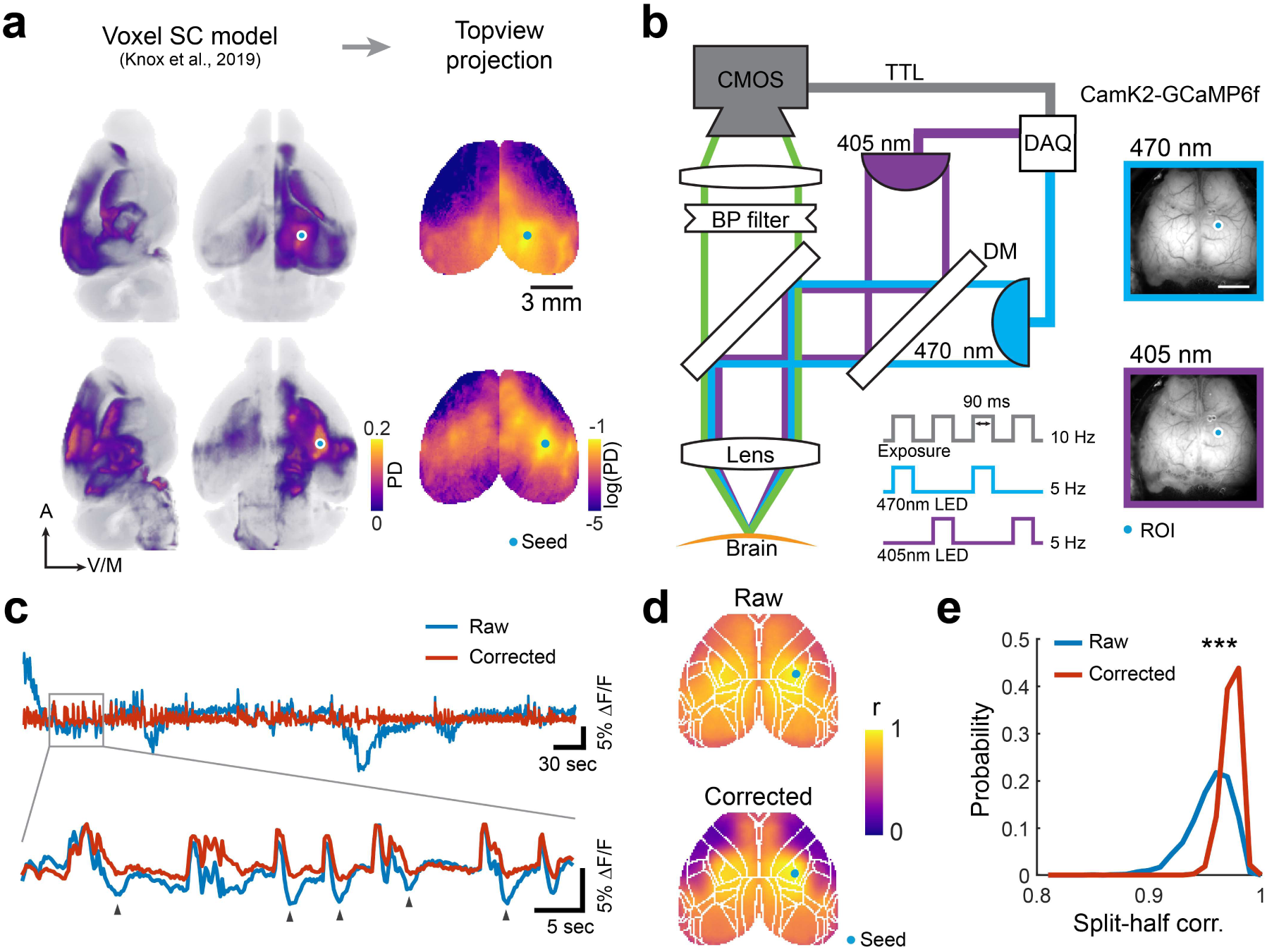
Mapping Structural and Functional Connectivity in Mouse Cortex. **a)** Representative voxel-level SC examples^34^. Each shows axonal projection density (PD) originating from a seed voxel (blue dot). The volumetric data are projected to topview maps and logarithmically scaled. **b)** Schematic of the dual-wavelength calcium imaging setup^35^. A 405 nm reference channel is used to correct calcium-independent artifact in the 470-nm GCaMP signal. **c)** Example traces of raw signal and corrected calcium-dependent signal extracted from the region of interest (ROI) indicated in **(b)**. Arrows indicate negative deflections characteristic of hemodynamic artifacts, which are attenuated in the corrected signal. **d)** Example functional connectivity (FC) maps from a seed (blue dot) derived from the raw and corrected signals. Note that the nonspecific global correlations, particularly between anterolateral and posteromedial areas observed in the raw data, are reduced in the corrected FC map. **e)** Assessment of cross-animal consistency via split-half correlation analysis. Histograms show the distribution of correlation coefficients between FC maps generated from random splits of the dataset (n = 14 mice; 3432 combinations). Signal correction significantly improved consistency between the dataset (raw r = 0.95 ± 0.02; corrected r = 0.97 ± 0.01; p < 0.001, Wilcoxon signed-rank test).

Resting-state FC was obtained by conducting wide-field calcium imaging in awake mice, which expressed GCaMP6f in excitatory neurons (Camk2-GCaMP6f, n = 14 mice). To reduce noise from non-neuronal components, we used a dual-wavelength imaging approach as previously described^35^ (**Fig. 1b**; see Methods). The raw GCaMP signal excited by 470 nm was corrected by a reference signal excited by 405 nm, which resulted in a reduced hemodynamic artifact^36^ (**Fig. 1c**). The FC from the corrected GCaMP signal showed enhanced contrast in reginal variations when compared to that obtained from raw signals excited by 470 nm (**Fig. 1d**). Furthermore, the signal correction significantly improved the consistency of the FC between subjects, as demonstrated by the result of split-half correlation (r = 0.97 ± 0.01 for corrected data and 0.95 ± 0.02 for raw data; Wilcoxon signed-rank test, p < 0.01; **Fig. 1e**).

We next examined the relationship between population-averaged FC and SC at whole-cortex level (**Fig. 2a** & **2b**). The logarithmically scaled SC and FC values were strongly correlated across the entire cortex (Spearman r = 0.72, R^2^ = 0.49; **Fig. 2c**), indicating that regions connected with denser axonal projections (high SC) exhibited synchronized calcium dynamics (high FC). However, this correspondence weakened in distal and contralateral areas, where high FC was observed despite low SC. Reflecting with this mismatch, a linear model based on SC accounted for only a limited fraction of FC variance (leave-one-out cross-validation R^2^ = 0.31 ± 0.14; **Fig. 2d**). Notably, SC showed stronger correlation with FC derived from the corrected GCaMP signal than with FC from the raw signal (paired t-test, p < 0.01; **Fig. 2c** & **2d**). Together, our approach combining viral tracing and calcium imaging revealed a strong yet incomplete S-F coupling in the mouse cortex^37^.

**Figure 2.**
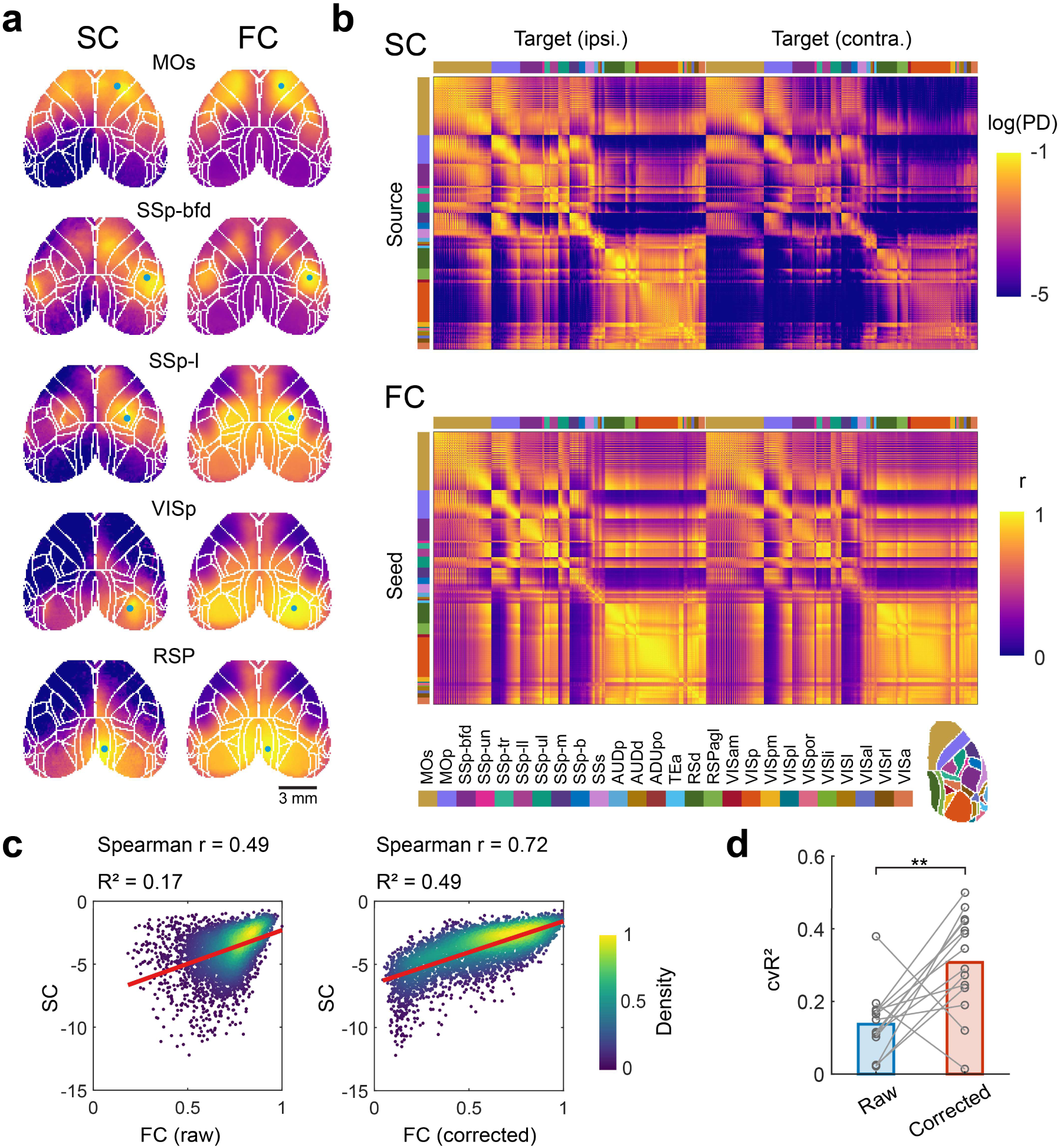
Comparison of Structural and Functional Connectivity in Mouse Cortex. **a)** Example SC and FC maps. Blue dots indicate the seed locations. The color scale corresponds to that used in (**b**). **b)** SC and FC connectivity matrices. Rows and columns are organized by 26 cortical areas defined by the Allen Mouse Brain Common Coordinate Framework. The SC matrix displays PD from sources to ipsilateral and contralateral targets; the FC follows an identical arrangement. **c)** Scatter plots comparing the log-scaled SC and FC values in all cortical regions. Each dot represents a single pixel and is color-coded by local density. Results using FC derived from both raw and corrected signals are shown. **d)** The cross-validated coefficient of determination (cvR^2^) of a linear model using SC to predict FC. Each point represents a one-fold validation. Signal correction shows a significant impact on the prediction accuracy (raw cvR^2^ = 0.14 ± 0.09; corrected cvR^2^ = 0.31 ± 0.14; p<0.01, paired t-test). Abbreviations: MOs, secondary motor; MOp, primary motor; SSp-bfd, primary somatosensory, barrel field; SSp-un, primary somatosensory, unassigned; SSp-tr, primary somatosensory, trunk; SSp-ll, primary somatosensory, lower limb; SSp-ul, primary somatosensory, upper limb; SSp-m, primary somatosensory, mouth; SSp-n, primary somatosensory, nose; SSs, supplemental somatosensory; AUDp, primary auditory; AUDd, dorsal auditory; AUDpo, posterior auditory; TEa, temporal association; RSPd, retrosplenial, dorsal; RSPagl, retrosplenial, lateral agranular; VISam, anteromedial visual; VISp, primary visual; VISpm, posteromedial visual; VISpl, posterolateral visual; VISpor, postrhinal visual; VISl, lateral visual; VISli, laterointermediate visual; VISal, anterolateral visual; VISrl, rostrolateral visual; VISa, anterior visual.

### Structure-Function Coupling Varied Systematically Across Cortex

The incomplete S-F coupling at the global level may stem from regional differences in the S-F relationship. To investigate this variation, we quantified local S-F coupling by calculating the Spearman correlation between the experimentally obtained FC and the SC-predicted FC at the pixel level (Fig. 3a). The SC-predicted FC was obtained using a linear regression model that fit SC to FC. The resulting cortical map revealed significant regional heterogeneity in S-F coupling (Fig. 3b). Although the spatial variation of S-F coupling was largely consistent between the ipsilateral and contralateral hemispheres (Pearson r = 0.89), the S-F coupling was weaker in the contralateral hemisphere. This overall reduction in S-F coupling in the contralateral hemisphere may reflect the contribution of indirect, polysynaptic connections to commissural FC^38^. We also observed similar S-F coupling maps across individual mice, confirming the robustness of these patterns (**Supplementary Fig. 3a**).

**Figure 3.**
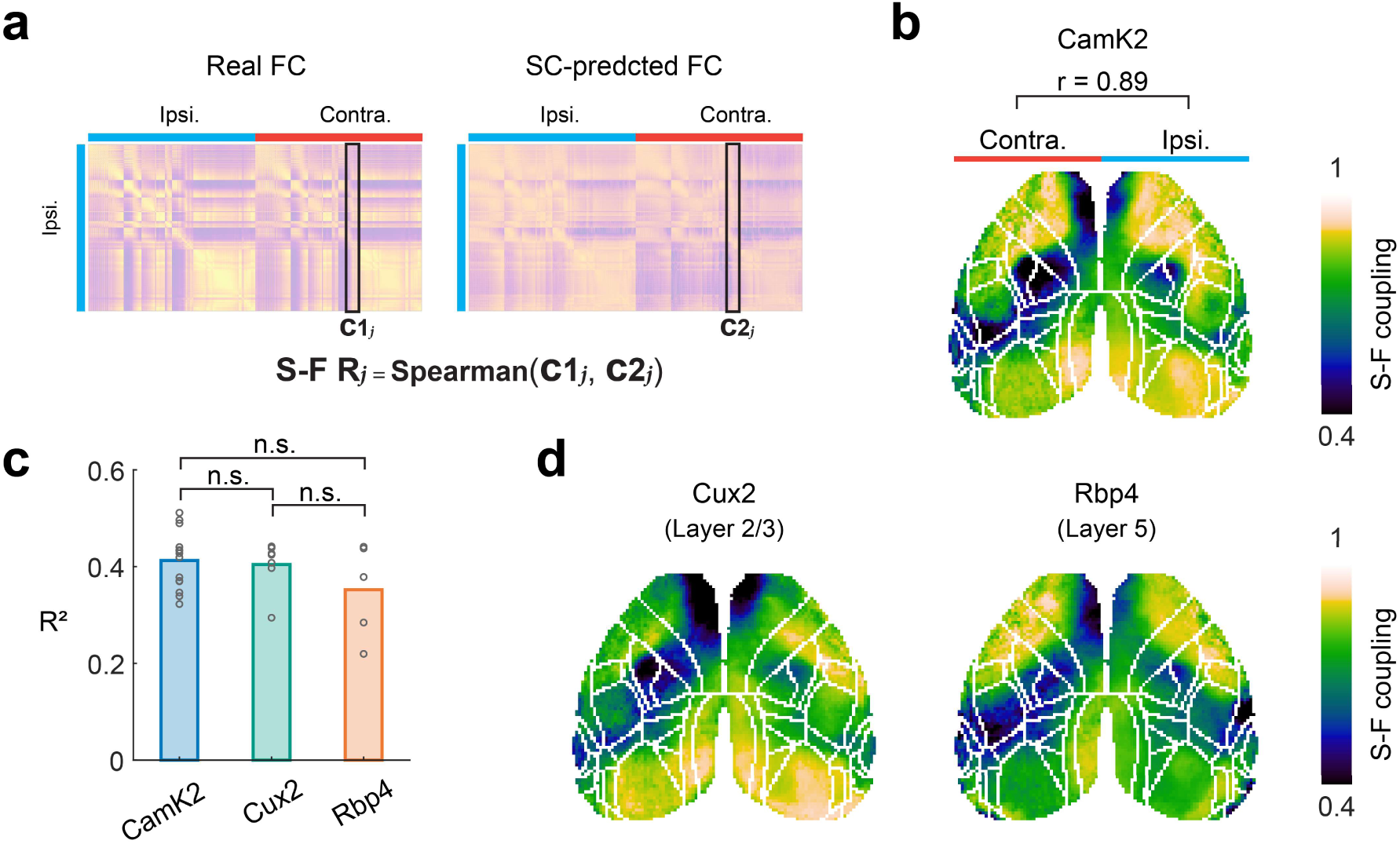
Heterogenous Structure-Function Coupling across the Cortex. **a)** Schematic illustrating the estimation of regional S-F coupling. Coupling strength was defined as the Spearman correlation between column vectors of the real FC matrix and the SC-predicted FC matrix. **b)** Cortical map of S-F coupling. The right hemisphere of the map displays ipsilateral S-F coupling, while the left hemisphere displays contralateral S-F coupling. The pearson correlation between them is shown. **c)** Global correspondence (R^2^) between SC and FC compared across three transgenic lines: Camk2 (full layer), Cux2 (layer 2/3), and Rbp4 (layer 5). Each point represents an individual mouse. Camk2 R^2^ = 0.41 ± 0.06; Cux2 R^2^ = 0.41 ± 0.05; Rbp4 R^2^ = 0.35 ± 0.09; p=0.23, one-way ANOVA. **d)** Cortical map of S-F coupling for the Cux2 and Rbp4 mice, demonstrating spatial patterns consistent with the Camk2 data shown in (**b**).

Intratelencephalic neurons in layer 2/3 and layer 5 primarily mediate corticocortical connections^29^. To test whether the observed regional differences in S-F coupling were unique to these layers or common across them, we conducted wide-field calcium imaging in two additional mouse lines: Cux2-GCaMP6f (n = 7) and Rbp4-GCaMP6f (n = 5), which express GCaMP in layer 2/3 and layer 5 pyramidal cells, respectively^39^. We repeated our comparison between the SC and layer-specific FC (**Supplementary Fig. 3c**) and found that both mouse lines exhibited FC patterns that were broadly similar to those of Camk2-GCaMP6f mice. The global correlation between layer-specific FC and SC did not differ significantly (one-way ANOVA, p = 0.23; Fig. 3c). Furthermore, the regional variation in S-F coupling was largely consistent across the different mouse lines (Fig. 3d & **Supplementary Fig. 3d**), indicating that any layer-specific differences in the S-F relationship were not evident. Because SC contains information on the direction of axonal projections, we also asked whether the SC direction affects S-F relationships. We found the spatial pattern of S-F coupling was not significantly affected by the directionality of the SC data (**Supplementary Fig. 3b** & **3c**). Taken together, these analyses demonstrate that regionally heterogeneous S-F coupling is a robust feature of the mouse cortex independent of cortical layers or direction of axonal projections.

### Structure-Function Coupling Covaried with Intrinsic Cellular Properties

The regional variation in S-F coupling suggests that FC is influenced by local cellular properties as well as SC. To better understand these cellular properties, we examined the relationship between regional S-F coupling and several potential factors, including myelination (estimated by T1w/T2w ratio and oligodendrocyte density), E/I balance, synaptic density, and spatial transcriptomic profiles.

We found that the regional S-F coupling was negatively correlated with two markers of myelination, including T1w/T2w ratio^30^ (Spearman r = −0.53, R² = 0.21; p = 0.01; Fig. 4a) and oligodendrocyte density^40^ (Spearman r = −0.63, R^2^ =0.21, p < 0.01; **Fig. 4b**). In addition, we observed significant negative correlations between the S-F coupling and the ratio of excitatory to inhibitory neurons^40^ (Spearman r = −0.66, R^2^ =0.29, p < 0.01; **Fig. 4c**). Consistently, the S-F coupling was also negatively correlated with the markers of excitatory synapses such as PSD95 (Postsynaptic Density 95; Spearman r = −0.57, R^2^ = 0.15, p = 0.02; **Fig. 4d**) and SAP102^41^ (Synapse-Associated Protein 102; Spearman r = −0.51, R^2^ = 0.18, p < 0.05; **Fig. 4e**). Therefore, cortical regions with higher myelination, a greater E/I ratio, and increased excitatory synapse density exhibited weaker S-F coupling. In contrast, S-F coupling did not show a significant relationship with the anatomical cortical hierarchy, which has been defined by the proportion of feedforward (FF) and feedback (FB) axonal inputs, based on the laminar distributions of axonal terminals^29^ (Spearman r = 0.16, R^2^ = 0.02, p = 0.44; **Fig. 4f**). For example, primary somatosensory areas (e.g., SSp-ll, SSp-bfd), which are low in the anatomically defined cortical hierarchy, exhibited lower S-F coupling (**Fig. 4f**). This lower S-F coupling in primary somatosensory areas was not attributed to spontaneous movements during the imaging (**Supplementary Fig. 4**).

**Figure 4.**
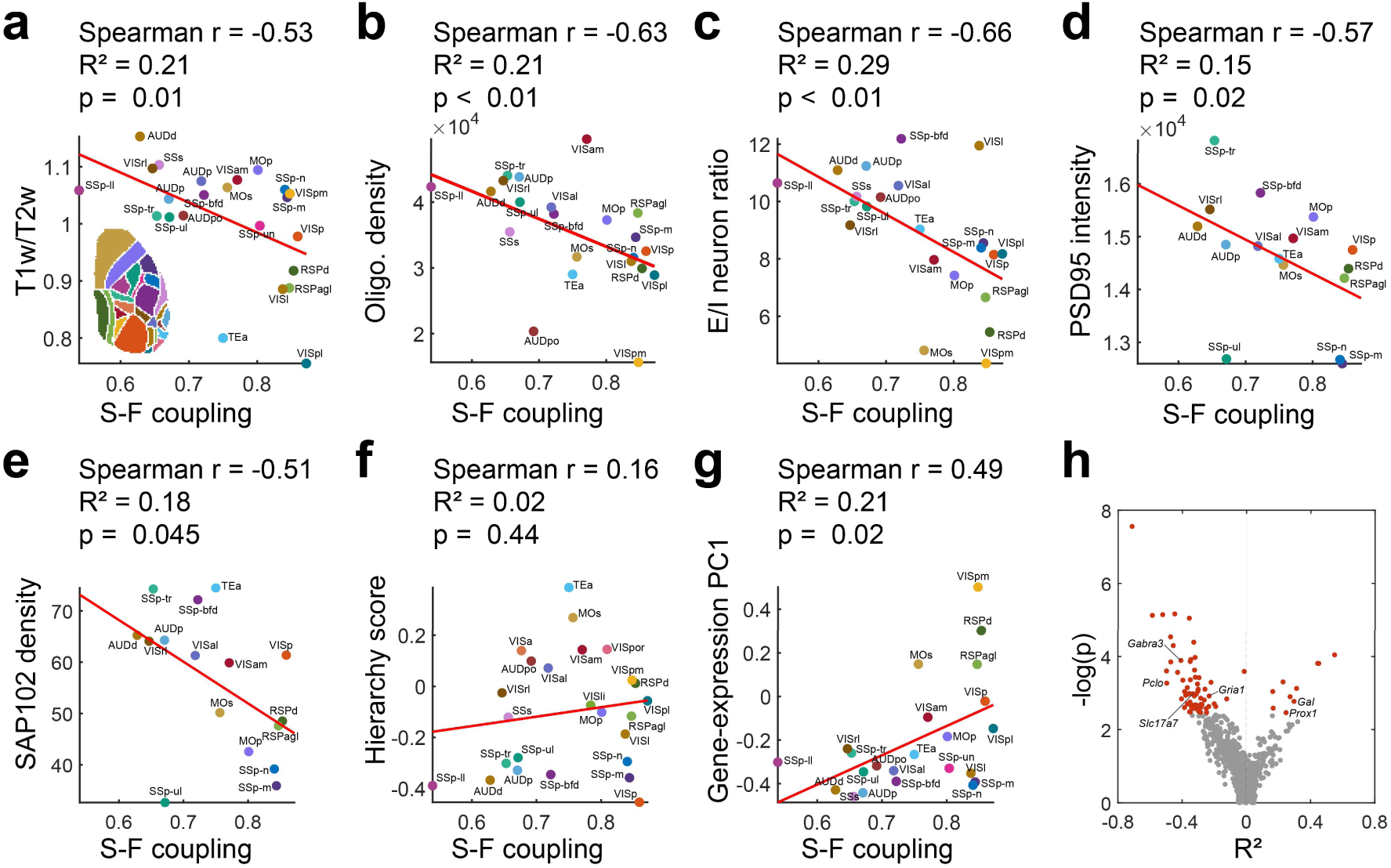
Structure-Function Coupling Covaried with Intrinsic Cellular Properties. **a)**-**g)** Scatter plots displaying the relationship between regional S-F coupling and intrinsic cellular properties: **a)** T1w/T2w ratio^30^; **b)** oligodendrocyte density^40^; **c)** ratio of excitatory and inhibitory (E/I) neuron density^40^; **d)** PSD-95 intensity (protein amount per area)^41^; **e)** SAP102 density (number of puncta per area)^41^; and **f)** hierarchy score^29^ defined by laminar FF/FB projections; and **g)** the first principle component (PC1) of 1,055 brain-expressing genes^30,42^. **h)** Volcano plot quantifying the association between individual gene expression and S-F coupling. Each point represents a single gene. The x-axis displays the R^2^ value signed by the direction of the correlation; the y-axis displays the uncorrected p-value (Spearman correlation). Red points indicate genes that reached statistical significance after False Discovery Rate (FDR) correction (q < 0.05). The full gene list can be found in **Supplementary Table 1.**

To investigate the molecular basis of these associations, we examined the relationship between the S-F coupling and the spatial expression pattern of 1,055 genes^30,42^. The first principal component (PC1) of these genes showed a significant correlation with the S-F coupling (Spearman r = 0.49, R^2^ = 0.21, p = 0.02; **Fig. 4g**). By computing the correlation coefficient for each gene, we identified 10 genes that were positively correlated and 64 genes that were negatively correlated with S-F coupling (FDR-corrected q < 0.05; **Fig. 4h** & **Supplementary Table 1**). Notably, two of the positively correlated genes were involved in inhibition of excitatory synaptic transmission: *Prox 1*, which regulates interneuron development^43^, and *Gal*, which encodes galanin, a neuropeptide known to inhibit glutamate release through G protein-coupled receptors^44^. In addition, the negatively correlated genes included multiple genes essential for the synaptic transmission, such as *Gria1*, encoding GluA1 glutamate ionotropic receptor subunit of AMPA receptors^45^, *Gabra3*, encoding gamma-aminobutyric acid (GABA) A receptor subunit alpha 3^46^, *Slc17a7*, encoding vesicular glutamate transporter 1 (VGLUT1)^47^, and *Pclo*, encoding presynaptic cytomatrix protein piccolo^48^. Taken together, these findings highlight the role of synaptic transmission in regulating S-F coupling. The overall trend suggests that stronger excitatory synaptic transmission combined with weaker inhibitory synaptic transmission, may lead to lower S-F coupling, particularly in the transmodal areas.

### Structure-Function Coupling Followed a Unimodal-to-Transmodal Gradient

In humans, regional variation in S-F coupling aligns with a principal functional gradient that captures hierarchical levels from unimodal to transmodal areas^16,22^. To examine whether the S-F coupling in mice exhibits a similar feature, we obtained the principal functional gradients from the FC in mice as described previously^18,49^. The first two gradients collectively accounted for 50% ± 4% of the variance (**Fig. 5a**). The first principal gradient (Gradient 1) spanned from the primary visual cortex at one end to the somatomotor areas at the other end and effectively separated areas for distinct sensory modalities. In contrast, the second principal gradient (Gradient 2) anchored unimodal sensorimotor areas at one extreme and transmodal default mode network regions at the other^50,51^ (**Fig. 5b**). We therefore interpreted Gradient 2 as reflecting the unimodal-to-transmodal gradient in the mouse cortex, which is homologous to the well-characterized principal functional gradient observed in humans^31,52^.

**Figure 5.**
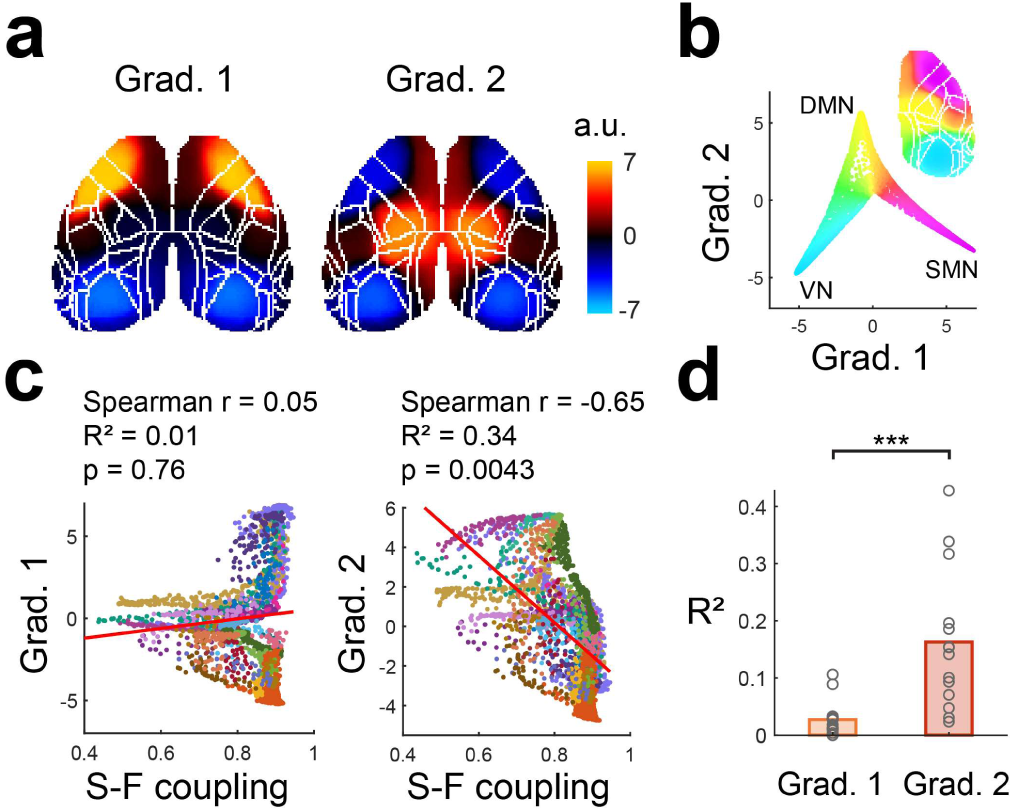
Structure-Function Coupling Followed a Unimodal-to-Transmodal Gradient. **a)** Cortical maps of the first two principal functional gradients derived from the population-averaged FC matrix. **b)** Gradient space representation. Scatter plot comparing Gradient 1 versus Gradient 2 values for each cortical pixel. Pixels are color-coded by their network affiliation, highlighting the segregation of three major networks: visual network (VN), somatomotor network (SMN), and default-mode network (DMN)^18^. **c)** Scatter plots comparing the S-F coupling against the scores of Gradients 1 and 2. Each dot represents a pixel on the cortex. Spatial correlations were assessed using the Moran permutation test. **d)** Bar plot shows the R^2^ for S-F coupling correlated against Gradients 1 and 2 for each mouse. (Gradient 1 R^2^ = 0.05 ± 0.04; Gradient 2 R^2^ = 0.25 ± 0.09; p < 0.001, paired t-test).

Consistent with this interpretation, the S-F coupling in mouse cortex correlated strongly with Gradient 2 (Spearman r = −0.65, R^2^ = 0.40; p < 0.01 in Moran permutation test) instead of Gradient 1 (Spearman r = 0.05, R^2^ = 0.01; p = 0.76; **Fig. 5c**). By analyzing the data of individual mice, we confirmed the S-F coupling specifically correlated with Gradient 2 and not Gradient 1 (parried t-test, p < 0.001; **Fig. 5d**). These findings demonstrated that the regional variation in S-F coupling in the mouse cortex followed a unimodal-to-transmodal gradient, recapitulating the patterns previously established in humans.

Our findings so far suggest that the basic principle of S-F coupling is conserved between humans and mice. To evaluate this objectively, we directly compared human S-F coupling data to the mouse S-F coupling data. We obtained human data from Collins et al.^53^, which quantified S-F coupling as the coefficient of determination (S-F R^2^; **Fig. 6a**). We therefore calculated the S-F R^2^ in mice from our datasets using the same approach. The spatial pattern of S-F R^2^ in mice was highly similar to that in our primary analysis (**Supplementary Fig. 6**). Next, we projected the human data onto the mouse brain using a recently developed tool that translates the data across species based on structural and transcriptional similarity^54^. The human values mapped to the mouse brain showed positive correlation with the mouse data (Spearman r = 0.49, R^2^ = 0.13, p = 0.02; **Fig. 6b**), indicating that the spatial topography of S-F coupling was evolutionarily conserved between the two species.

**Figure 6.**
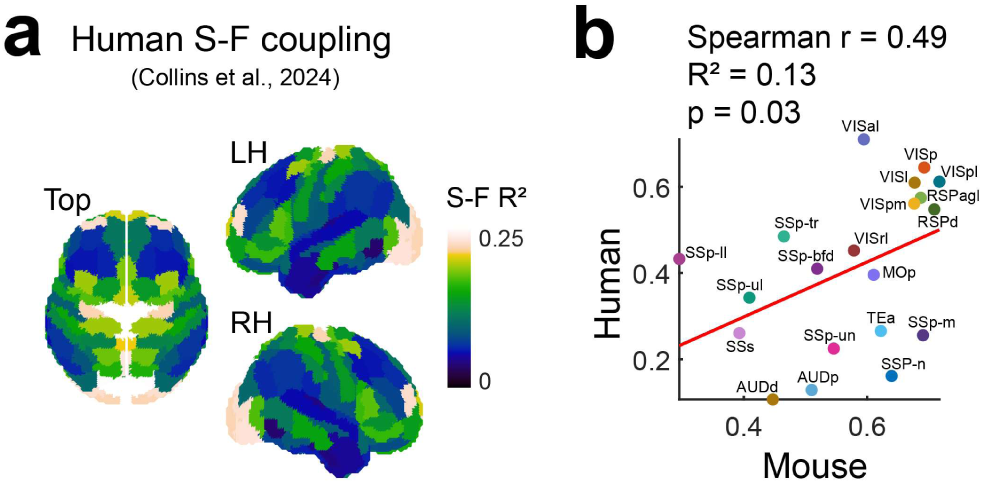
Cross-Species Conservation of Structure-Function Coupling. **a)** Regional S-F coupling in human cortex^53^. Coupling strength is quantified as the coefficient of determination (S-F R^2^) derived from linear models using SC to predict FC. **b)** Cross-species comparison of regional S-F coupling. Human S-F R^2^ data were mapped onto homologous mouse cortical areas using a cross-species translation framework^54^ and compared against mouse S-F R^2^ data.

## Discussion

In this study, we examined the relationship between SC and FC in mice. We used axonal tracing data to assess SC and wide-field calcium imaging to obtain FC at a fine spatial resolution. Our findings revealed that axonal SC does not completely account for FC, and S-F coupling varies across different regions of the mouse cortex. We discovered that regional differences in S-F coupling were correlated with several intrinsic cellular properties, including myelination, E/I neuron ratio, excitatory synaptic density, and spatial transcriptomic patterns, but not with the anatomically defined cortical hierarchy. Notably, we demonstrate that the S-F coupling in mice follows a principal functional gradient that extends from unimodal sensory areas to transmodal association areas, which recapitulates the observation in the human cortex. Taken together, these findings establish the fundamental properties of the S-F relationship that are conserved between humans and mice, providing a framework for future studies aimed at clarifying cellular and molecular mechanisms that regulate mesoscale cortical dynamics in both health and disease.

The spatial organization of S-F coupling aligns with a unimodal-to-transmodal functional gradient, a feature conserved across mammals^52^. The S-F coupling is stronger in the unimodal cortex, meaning that the anatomical connectivity provides a “hard” architecture optimized for efficient information transmission. In contrast, the S-F coupling is weaker in the transmodal areas, suggesting that these areas have a “soft” architecture mediated by uniquely refined microstructural and molecular properties that facilitate flexible computation necessary for context-dependent cognition^21^. Mechanistically, our results suggest this flexibility is driven by an elevated E/I balance (increased density of excitatory neurons and synapses and stronger excitatory synaptic transmission). Strong local recurrent excitation is known to amplify intrinsic neural reverberation, thereby allowing local circuit dynamics to override the rigid constraints of anatomical inputs. By reducing the dependence of activity on structure, the elevated E/I balance may serve as a mechanism for a reservoir of internal states and expand the timescale of information integration^55,56^.

While the functional gradient of S-F coupling was conserved in mice and humans, its relationship to two anatomical markers showed notable divergences between mice and humans. First, we identified a significant difference in the relation to myelination. In the mouse cortex, areas with higher myelination showed weaker S-F coupling (**Fig. 4a & 4b**). This relationship contrasts with findings in humans, where greater myelination is associated with a stronger S-F coupling^22^. Typically, myelination is thought to enhance a ‘hard-wired’ circuitry by increasing transmission speed and restricting plasticity^57^. Although this model aligns with the characteristics of the human unimodal cortex, it does not match our observation in mice. One plausible explanation is that myelination in rodents may not follow the organizational principles observed in primates, given that the mouse cortex exhibits less distinct laminar and myeloarchitectural differentiation than that of primates^29,30^. Second, contrary to our expectation, S-F coupling did not correlate with the anatomically defined hierarchy based on feedforward/feedback (FF/FB) axonal projections^29^. While this is counterintuitive, we interpret this result with caution. The anatomically defined hierarchical score showed significant correlation with some intrinsic cellular properties—such as the T1w/T2w ratio and the gene expression PC1 (data not shown). Because these properties correlated with S-F coupling, the lack of significant correlation between S-F coupling and anatomical hierarchy may be due to insufficient statistical power.

Compared with humans, S-F coupling in the mouse cortex was generally stronger and more spatially uniform. This pattern mirrors findings in macaques^58^, suggesting a cross-species trend in which larger and more complex brains exhibit a weaker anatomical constraint and greater functional flexibility. A plausible driver of this shift is the evolutionary expansion of transmodal association areas in primates^59^. Consistent with this hypothesis, the unimodal-to-transmodal gradient typically emerges as the *principal* functional gradient in primates^18,19^, whereas it appears as a *secondary* gradient in mice ^25,32^ (**Fig. 5a**) and in zebrafish^60^. Instead, the principal functional gradient in both mice and zebrafish follows a visuomotor axis. The disproportionate expansion of the transmodal cortex represents a defining feature of primate evolution and likely contributes to the progressive decoupling of function from structure along the mammalian lineage.

In summary, our study identifies a conserved principle of cortical organization: a graded decoupling of function from structure that occurs along the unimodal-to-transmodal cortical gradient. By validating this gradient in mice, we bridge a critical gap between human neuroimaging—which has long characterized these patterns but lacks cellular resolution—and rodent circuitry, which offers mechanistic insights. The alignment of this macroscopic gradient with microscopic properties positions the mouse as a valuable translational model. This allows us to use targeted genetic manipulations and longitudinal imaging to dissect the precise cellular, molecular, and developmental mechanisms that influence the relationship between brain structure and function.

## Materials and Methods

### Animals

All animal experiments were approved by and performed under the guidelines of the RIKEN Animal Experiment Committee. The imaging experiments were conducted on adult (9-23 weeks old) male and female mice generated by crossing CaMKIIα-Cre (RRID: IMSR_JAX:005359, from Jackson Laboratory), Cux2-Cre (RRID: MMRRC_031778-MU, from Mutant Mouse Resource and Research Center at University of Missouri), or Rbp4-Cre (RRID: MMRRC_037128-UCD, from Mutant Mouse Resource and Research Center at University of California at Davis) lines^39^ with Ai95 (RCL-GCaMP6f)-D (RRID: IMSR_JAX:024105, from Jackson Laboratory).

### Surgery

Surgery was performed to implant a head plate on the skull using adapted procedures described previously^35,61^. Briefly, the anesthesia was induced with 5% and maintained with 1-2% isoflurane. Following scalp dissection, a custom-made stainless head plate was attached to the skull using Super-Bond C&B dental cement (Sun Medical Co., Ltd.). A thin layer of UV-curable nail resin (Top Gel Glossy; Jelly Nail) was applied onto the skull to make it optically transparent.

### Wide-field Calcium Imaging

Dual-wavelength wide-field calcium imaging and data preprocessing were performed as described previously^35^. The head-fixed awake mice were imaged in darkness using a macro zoom fluorescence microscope (MVX10; EVIDENT) equipped with a 1× objective (1× MVX Plan Apochromat Lens, N.A. 0.25) and an sCMOS camera (Andor Zyla 4.2 PLUS; Oxford Instruments). The camera exposure time was set to 100 ms, resulting in a 5-Hz sampling rate per channel. Alternating 405-nm LED (M405L4; Thorlabs, Inc.) or 470-nm LED (M470L5; Thorlabs, Inc.) was triggered for 80 ms during the imaging. A shield was attached to the headplate to protect the eye from illumination light.

### Calcium Imaging Data Preprocessing

All data processing and analyses were performed using MATLAB (R2023a, MathWorks, Inc.) following established protocol^35^. Briefly, motion-corrected frames were downsampled to 256 × 256 pixels. For each channel, data were normalized to ΔF/F by subtracting and then dividing by the mean fluorescence across all frames. The resulting 470-nm ΔF/F signal was used as the raw GCaMP signal. To correct the raw signal, the 470-nm ΔF/F signal was first denoised using singular value decomposition (SVD) and temporally filtered with a 0.1 Hz high-pass filter (4th-order Butterworth). The 405-nm reference ΔF/F signal was then regressed out from the 470-nm ΔF/F signal. The resulting residual data were used as the corrected GCaMP signal.

### Structural Connectivity Reconstruction

Voxel-level SC in the mouse cortex was reconstructed using a model developed by Knox et al.^34^, which is based on axonal tracing data in AMBCA^33^. The model predicts PD based on the location of the virus injection site. To construct a comprehensive cortical SC matrix, we simulated a dense array of virtual injections. Each virtual injection consisted of a one-voxel-wide path traversing through the cortical depth. The injection density along this path was modeled as a Gaussian distribution (µ = 0, σ = 0.3 mm) to mimic the superficial bias of wide-field imaging (**Supplementary Fig. 1b**). Model outputs were projected into a 2D space by first summing values along the orthogonal paths to the cortical surface and then mapping them to a horizontal plane^62^ (https://github.com/AllenInstitute/ccf_streamlines; **Supplementary Fig. 1c**). Because AMBCA viral injections were restricted to the right hemisphere, this procedure generated 3,518 SC maps, each estimating the PD distribution from a single pixel seed in the ipsilateral (right) hemisphere. We excluded maps with seeds located on the area borders, and the remaining 2,712 maps were vectorized and visualized as the SC matrix in **Fig. 2b**.

### Functional Connectivity Reconstruction

Imaging data were co-registered to the AMBCA reference space (Allen Mouse Brain Common Coordinate Framework, CCF^62^) via affine transformation using five manually selected skull landmarks (left and right ends of frontal pole, front center, lambda, and bregma). The data were further downsampled to 114 × 114 pixels to match the resolution of SC data. The FC matrix corresponding to the SC matrix was obtained by calculating the Pearson correlation of the ΔF/F activity between the 2,712 seed pixels in the right hemisphere and all 5,420 pixels in both hemispheres. The FC data were averaged first across two 20-min imaging sessions per mouse and then across different mice to yield the population-averaged FC matrix in **Fig. 2b**. We assessed the across-animal consistency of FC by splitting the mice into two halves (a total of 3,432 combinations for 14 mice) and computing the correlation between the resulting group-averaged FC in **Fig. 1e**.

### Comparison Between Structural and Functional Connectivity and Estimation of S-F Coupling

Spearman correlation was calculated between the SC and FC matrices to assess their correspondence at the whole-cortex level. We further quantified the SC’s predictive power using a linear regression model with leave-one-out cross-validation. In each fold, the model was trained on the mean FC map of n-1 animals (using SC as the predictor) and tested on the held-out animal. The cvR² was computed as the R² between predicted and observed FC for the held-out animal. To quantify the local S-F coupling, we calculated the Spearman correlation between corresponding column vectors of the population-averaged FC and SC-predicted FC matrices. The SC-predicted FC was obtained from a linear regression model that fitted FC from SC. Of note, this calculation can also be done row-wisely (**Supplementary Fig. 3b**), resulting in cortical maps that were highly similar to those from the column-wise calculation (Pearson r > 0.8; **Supplementary Fig. 3c**).

### Associating S-F coupling to Intrinsic Cellular Properties

We correlated the regional S-F coupling with a set of cellular properties across 26 cortical areas. Regional S-F coupling values were obtained by averaging all pixels within the region in the right hemisphere. Due to an update of the CCF parcellation, several areas were replaced by new parcels. We matched PTLp (posterior parietal association areas) to VISrl and treated other areas (VISa, VISli, and VISpor) as missing data. T1w/T2w ratio was obtained from Fulcher et al.^30^ and can be found at https://figshare.com/articles/dataset/Mouse_cortical_gradients/7775684. Anatomical hierarchy score based on the laminar projection pattern of axonal terminals is from Harries et al.^29^; excitatory and inhibitory neuron densities, as well as oligodendrocyte density, are from Erö et al.^40^; PSD95 and SAP102 density and intensity data were obtained from Zhu et al.^41^. Transcriptomic analysis utilized processed gene-expression data from Fulcher et al.^30^. Briefly, the original gene-expression data were obtained from AMBA (Allen Mouse Brain Atlas)^42^ and filtered by quality control and brain enrichment, yielding expression levels of 1,055 genes. Probabilistic PCA was then performed to extract the dominant expression patterns.

### Principal Gradient Analysis

Principal functional gradients were extracted from population-averaged FC using the BrainSpace toolbox^49^. We employed the PCA approach with a normalized angle kernel. To test the significance of spatial correlation between gradient and S-F coupling maps, we used a spatial permutation test based on Moran spectral randomization in the BrainSpace toolbox to construct a null distribution that preserves spatial autocorrelation. All tests were performed on the right hemisphere. Individual-level analysis was performed by comparing the gradients and S-F coupling derived from individual animal data.

### Human-to-Mouse Brian Translation

We used TransBrain toolbox^54^ (https://github.com/ibpshangzhe ng/Transbrain) to map human S-F R^2^ values onto the mouse cortex. The S-F R^2^ data were obtained from a previous human study^53^. The data were originally parcellated in the Yale Brain Atlas (https://yalebrainatlas.github.io/YaleBrainAtlas/download.html), remapped to the Brainnectome Atlas (https://atlas.brainnetome.org/download.html), and then translated to mouse data in CCF parcellation using TransBrian. Some areas, such as MOs and VISam, were missing in this translation. We correlated the remapped human values with mouse S-F coupling data (S-F R^2^) across 21 matched cortical areas.

## Data and Code Availability

The original AMBCA data are available at http://connectivity.brain-map.org. The voxel SC model can be obtained from https://github.com/AllenInstitute/mouse_connectivity_models. The code and data are available upon request to the corresponding authors.

## Conflicts of Interests

The authors have no competing conflicts of interests to declare.

## Author Contributions

Conceptualization: RL, AI & TM. Data acquisition: MT, MK, AI & RL. Data analysis: RL & TM. Manuscription writing: RL, TM & AI.

## Acknowledgements

We thank other lab members for their help. This work was supported by the following grants: JSPS KAKENHI (25K1856 to RL, 24H02331 to TM, and 22H05158 to AI); JST-CREST (JPMJCR22N4 to TM); AMED (JP25wm0625422 to TM).

## Notes

### Competing Interest Statement

The authors have declared no competing interest.

