## Supplementary Information for "Structure-Function Coupling Aligns with a Unimodal-to-Transmodal Gradient in the Mouse Cortex"

†Corresponding authors:

This file includes:

**Supplementary Materials**

**Supplementary Figures 1-6**

**Supplementary references**

### Supplementary Materials

#### SC Directionality and S-F Coupling

Unlike SC derived from diffusion MRI, SC based on axonal tracing is directional. *Output* SC represents axonal projection targets from the injection sites of viral tracers. In contrast, *input* SC shows the distribution of regions projecting to a given area. We therefore calculated the S-F coupling in two complementary ways. We calculated source S-F coupling for pixels (rows) in the ipsilateral hemisphere to capture the correlation between the FC and the *output* SC. Likewise, we calculated target S-F coupling for pixels (columns) in both hemispheres to capture the correlation between the FC and *input* SC (**Supplementary Fig. 3b**). For simplicity, we only used target S-F coupling in our main text.

The analysis using the source and target S-F coupling was generally consistent. For example, the source and target S-F coupling were strongly correlated in the ipsilateral hemisphere (Pearson r = 0.84, $R^2 = 0.71$), indicating that the regional S-F coupling is largely unaffected by the SC direction (**Supplementary Fig. 3c**). In addition, both source and target S-F coupling aligned with the same principal component of the FC. The source S-F coupling was significantly aligned with Gradient 2 (Spearman r = -0.65, $R^2 = 0.40$; $p < 0.01$ in Moran permutation test) instead of Gradient 1 (Spearman r = 0.16, $R^2 = 0.02$; p = 0.42 in Moran permutation test). However, we noticed more difference between the source and target S-F coupling after we summarized the whole-cortex data to 26 cortical areas (Pearson r = 0.73, $R^2 = 0.51$) and associated the results with intrinsic cortical properties. For example, the source S-F coupling showed a significant correlation with oligodendrocyte density (Spearman r = -0.45, $R^2 = 0.16$, $p < 0.05$) but not with T1w/T2w ratio (Spearman r = -0.38, $R^2 = 0.05$, p = 0.08). In addition, the source S-F coupling showed negative correlation with PSD95 intensity (Spearman r = -0.51, $R^2 = 0.12$, $p < 0.05$) but not with SAP102 density (Spearman r = -0.19, $R^2 = 0.06$, p = 0.47). Because the overall trend was consistent across source and target S-F coupling, we used only target S-F coupling in our main text. In addition to noise in the data, we speculate that differences between source and target S-F coupling can be due to methodological bias in viral tracing. The axonal PD near the injection sites is often overestimated due to contamination from infected somas, and source S-F coupling may have been more sensitive to this bias.

#### Spontaneous Movements and S-F Coupling

To estimate the potential influence of spontaneous movement on the S-F coupling in mice, we recorded the orofacial movements and pupil size during the macroscopic imaging. The faces of the mice were illuminated by an 850-nm infrared LED ring light (LED-R850IR; Arms System Co., Ltd.) and recorded by a high-speed CCD camera (HAS-U1; DITECT Co., Ltd.) that was synchronized with the calcium imaging camera (Supplementary Fig. 4a). We used DeepLabCut[1] (version 2.3.5) to trace the orofacial movements from the recorded videos. Specifically, we labeled 13 landmark points (4 surround the eye, 3 on the nose, 3 on the whisker pad, and 2 on the mouth; **Supplementary Fig. 4a**)[2] on 120 frames taken from 6 mice to train a ResNet-50-based model with default parameters for 20,000 iterations (test error = 7.46 pixels, the frame size was 1280 × 1024). The outlier points (points diverged largely from the local median) and the points with a likelihood of less than 0.5 were removed from the model-predicted traces. The missing points were filled by modified Akima cubic interpolation. Finally, we calculated the pixel distance between each pair of points and applied z-score normalization (**Supplementary Fig. 4b**).

To segment pupil area from the recorded videos, we used a U-net-based model (https://github.com/milesial/Pytorch-UNet). Specifically, we cropped the video around the eye and annotated the pupil area in 300 frames taken from 6 mice to train the model with 50 epochs (validation Dice score = 0.94). We then measured the pupil diameter by first fitting the annotated areas to an ellipse and averaging the ellipse's major and minor axes. We smoothed the pupil diameter traces to remove the outliers.

To quantify the impact of orofacial movements and pupil changes on the FC, we used a ridge regression model[3] . Briefly, we concatenated the face movement and pupil diameter data to predict the first 100 SVD temporal components of the calcium imaging data. Using a tenfold cross-validation, we computed the explained variance (cv$R^2$) of the prediction, which was projected back to the cortical space (**Supplementary Fig. 4d**). This gave us an estimation of the movement-predictable calcium activity. Then, we calculated the residual calcium activity by subtracting the movement-predictable calcium activity from the full calcium activity. Using this residual calcium activity, we calculated FC as the residual FC and compared it with the SC using a linear model.

While the residual FC showed a slight but significant increase in its global coupling with SC (**Supplementary Fig. 4f**), the local S-F coupling map remained virtually unchanged from the original data (Pearson r = 0.99; **Supplementary Fig. 4g**). The consistency in S-F coupling derived from the raw FC and the residual FC indicated that the S-F coupling at rest is largely unaffected by the spontaneous orofacial movements or changes in the arousal level during imaging.

**Robustness Analysis on S-F coupling**

To ensure the robustness of our findings against methodological variations, we employed another widely used method to quantify the S-F coupling[4]. Specifically, we used a multiple linear regression model to predict the functional connectivity profile of each pixel. The regressions were performed using SC as the only predictor (SC-only) or in conjunction with a set of geometric and SC-derived profiles (SC-augmented; **Supplementary Fig. 6c**). The additional predictors included the Euclidean distance between pixels on the brain, the path length between nodes (pixels), and communicability between nodes[5] . We calculated path length and communicability by following the previous descriptions[4] .

The S-F $R^2$ and our primary calculation of S-F coupling are two closely relevant metrices with a strong correlation (Pearson $r = 0.92$, $R^2 = 0.85$; **Supplementary Fig. 6a** & **6b**). Additional predictors improved both global SC-FC correspondence (**Supplementary Fig. 6d**) and local S-F coupling, especially within contralateral hemispheres (**Supplementary Fig. 6e**). S-F $R^2$ resulted from SC-only was used in **Fig. 6**.

**Statistical Analysis**

All statistical analyses were performed using MATLAB (R2023a, MathWorks, Inc.). The statistical significance threshold was set to 0.05.

*Split-Half Correlation of FC*

To assess the variation of FC between biological replicates, we randomly divided the data from 14 mice into two equal groups ($n = 7$ each). For each group, we calculated the population-averaged FC maps using both the raw and the corrected calcium signals and obtained Pearson correlation coefficients between the two groups. We repeated this process for all 3,432 unique combinations of how the mice were grouped. We compared the distributions of correlation coefficients between the raw and corrected signals using a two-sided Wilcoxon signed-rank test (**Fig. 1e**).

*Global SC-FC Correlation Analysis*

The global correspondence between SC and FC was quantified using a linear regression model where the SC was used to predict the FC. We assessed the model performance by calculating the coefficient of determination ($R^2$). To ensure robustness, a leave-one-out cross-validation scheme was employed, where the model was trained on data from all but one mouse and tested on the held-out mouse and was repeated for all mice. The resulting $R^2$ values for models using raw versus corrected GCaMP signals were compared using a paired t-test (**Fig. 2d**). Group comparisons of $R^2$ values across the three transgenic mouse lines were performed using a one-way ANOVA (**Fig. 3c**).

*Spatial Correlation and Permutation Testing*

To test the significance of correlations between two spatial maps (e.g., local S-F coupling and a functional gradient), we used a null model that accounts for spatial autocorrelation, which violates the independence assumption of standard correlation tests. We employed Moran spectral randomization[6,7] to generate a surrogate null distribution (20,000 permutations) of Spearman correlation coefficients. This method shuffles the map values while preserving their spatial structure. An empirical value was then calculated by comparing the true correlation coefficient to this null distribution. All tests were performed in the right hemisphere.

### Supplementary Figures

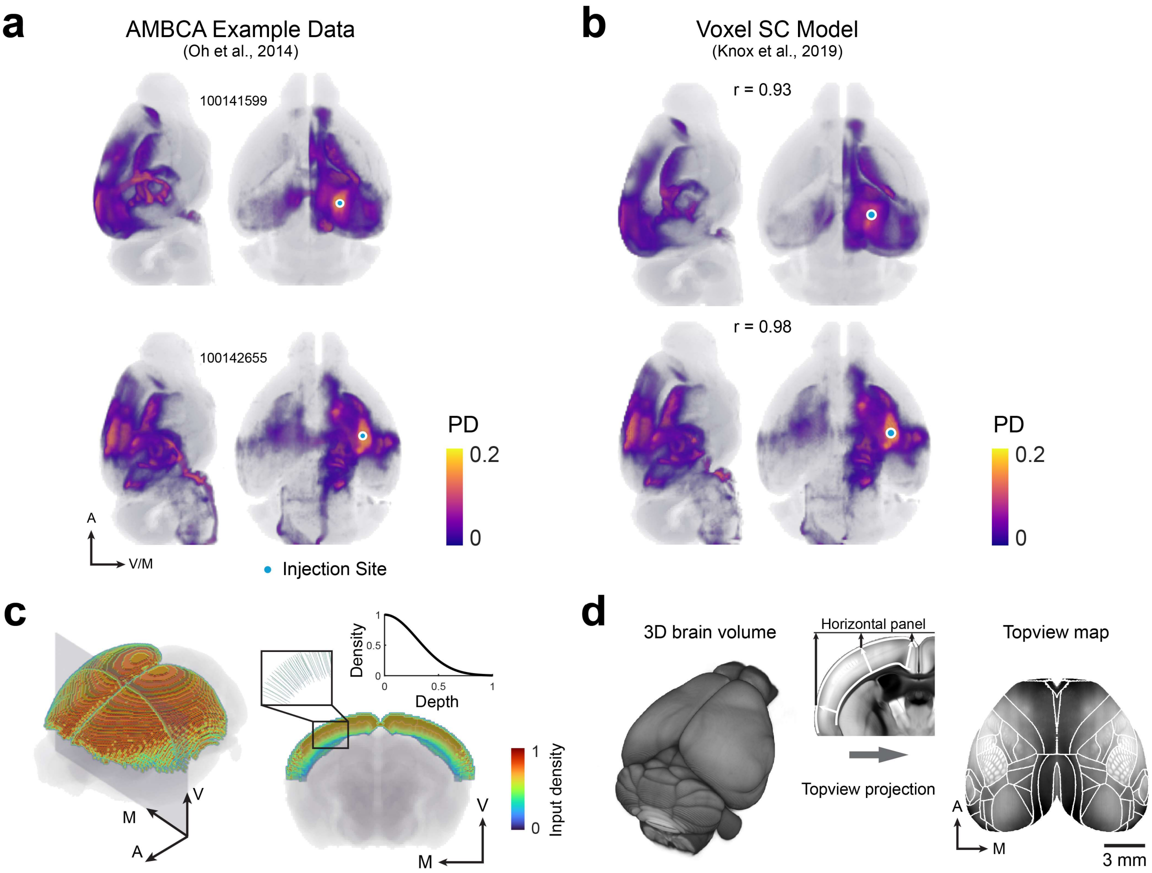


#### Supplementary Fig. 1. Reconstructing Structural Connectivity from AMBCA Data.

**a)** & **b)** Two examples of AMBCA data[8] . The injection site information from these experiments was used as input for the voxel-level SC model to generate the predictions shown in **b)** replotted from **Fig. 1a**. The AMBCA experiment ID and the Pearson r between the real data and the model prediction are given. **c)** To construct a comprehensive SC matrix in the cortex, we simulated a dense array of virtual injections. Each virtual injection consisted of a one-voxel-wide path traversing the cortical depth. The simulated injection density along this path followed a Gaussian distribution ($\mu = 0$, $\sigma = 0.3$ mm) to mimic the bias of wide-field optical imaging towards superficial layers. **d)** A schematic of the method used to project 3D cortical volumes into 2D top-down maps[9] . First, the volumetric cortical PD was averaged along paths perpendicular to the cortical surface. Second, these averaged values were projected onto a 2D plane. The image shown is an example of this projection method applied to a 3D autofluorescence volume from the Allen Institute.

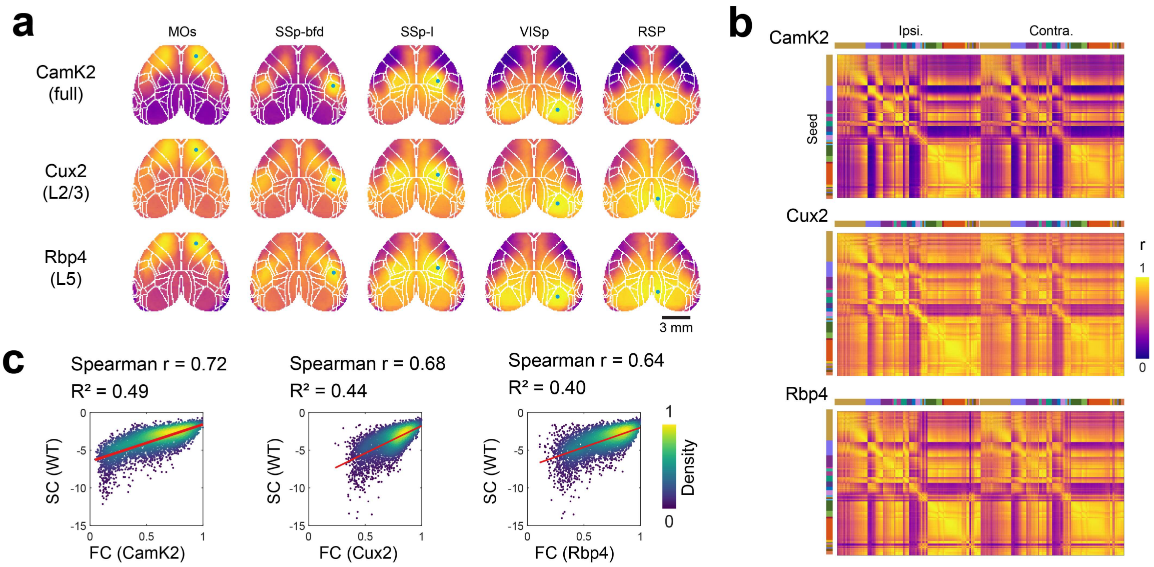


**Supplementary Fig. 2. Functional Connectivity in Different Cortical Layers.**

**a)** Example FC maps from three transgenic mouse lines with different layer-specific expression patterns. Top row: FC from CamK2-GCaMP6f mice (cortex-wide expression), reproduced from **Fig. 2a**. Middle and bottom rows: FC from Cux2- (layer 2/3) and Rbp4-GCaMP6f (layer 5) mice, respectively. The same color scale is used in (**a**) and (**b**). **b)** Similar to **Fig. 2b**. Pixel-wise FC matrices for the three transgenic lines. **c)** Similar to **Fig. 2c**. The global correlation between SC and FC for each of the three transgenic lines.

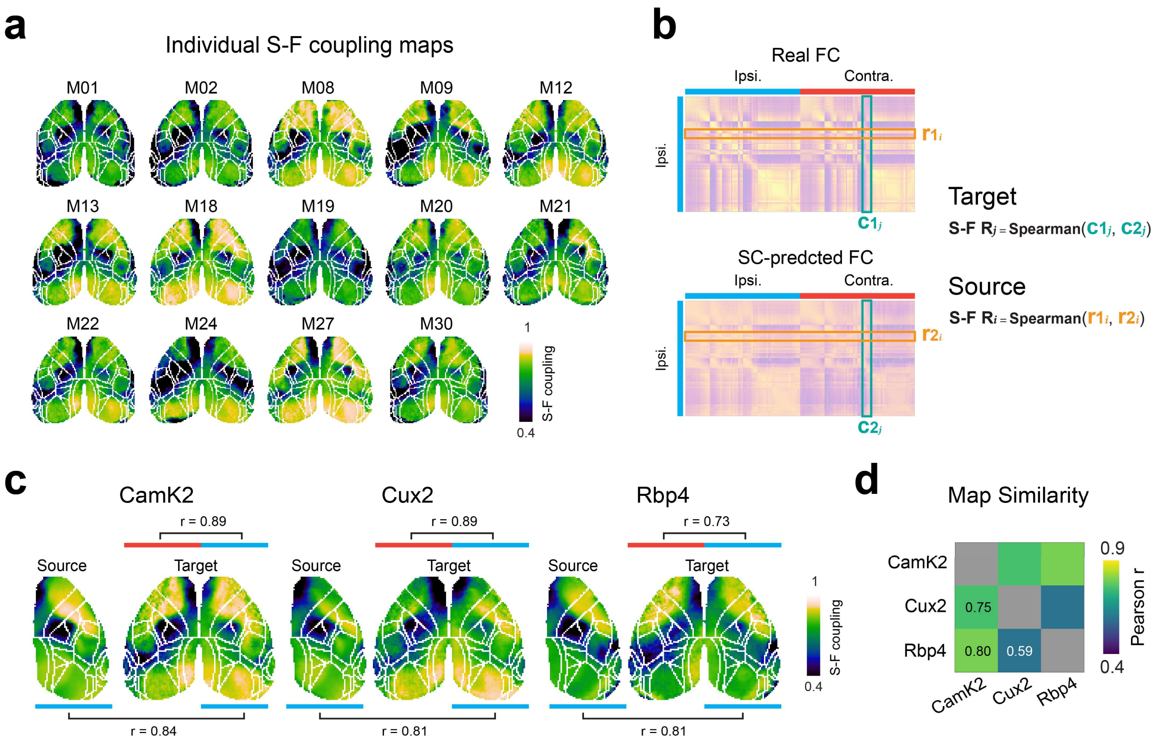


**Supplementary Fig. 3. Heterogeneous Structure-Function Coupling across the Cortex.**

**a)** Cortical maps of local S-F coupling for all 14 individual mice. **b)** In regarding the SC directionality, local S-F coupling can be calculated in two complementary ways: source and target maps were obtained by calculating the Spearman correlation of row and column vectors between the real and SC-predicted FC matrices, respectively. We used target S-F coupling in our main text. **c)** S-F coupling maps of three transgenic lines. Both source and target maps are shown. All maps revealed a consistent regional heterogeneity, indicating the local S-F coupling is insensitive to the direction of axon projection. **e)** The similarity (Pearson r) between the target S-F coupling maps of the three transgenic lines, demonstrating a high degree of consistency in the S-F relationship across different cortical layers.

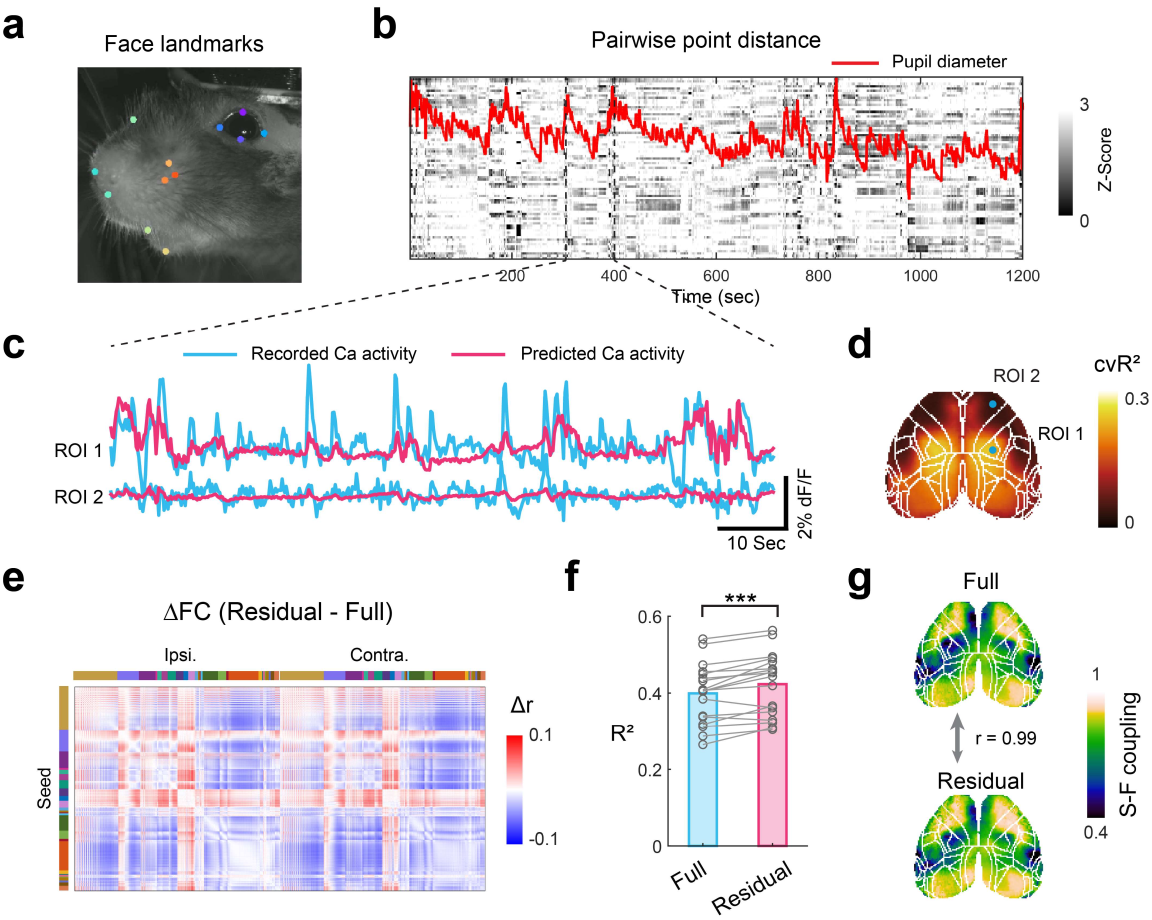


**Supplementary Fig. 4. Spontaneous Orofacial Movements Do Not Account for the Local Variation in S-F Coupling.**

**a)** An example video frame of a mouse's face, recorded concurrently with wide-field calcium imaging. Colored dots indicate facial landmarks tracked with DeepLabCut[1] . **b)** An example raster plot of orofacial movement from a single recording, quantified as the z-scored pair-wise distances between the tracked landmarks. The red trace represents changes in the pupil diameter. The gray lines indicate the time window in (**c**). **c)** Example traces of recorded calcium activity and the activity predicted from orofacial movements (red) using a ridge regression model[3] . **d)** A map of cross-validated R-squared (cv$R^2$) values, indicating the proportion of variance in local calcium activity that can be explained by spontaneous orofacial movements. **e)** The change in FC after regressing out movement-related components from the calcium signals (the difference between the original full FC and the residual FC after removing the movement-related components). **f)** The change in the global SC-FC correlation after removing movement-related components. The R-squared values are shown for both the full FC and the residual FC. Each dot represents an individual mouse. The improvement is small but significant ($p < 0.001$, paired t-test). **g)** A comparison of the S-F coupling maps calculated from the full FC (left) and the residual FC after movement correction (right), demonstrating high similarity (Pearson $r = 0.99$).

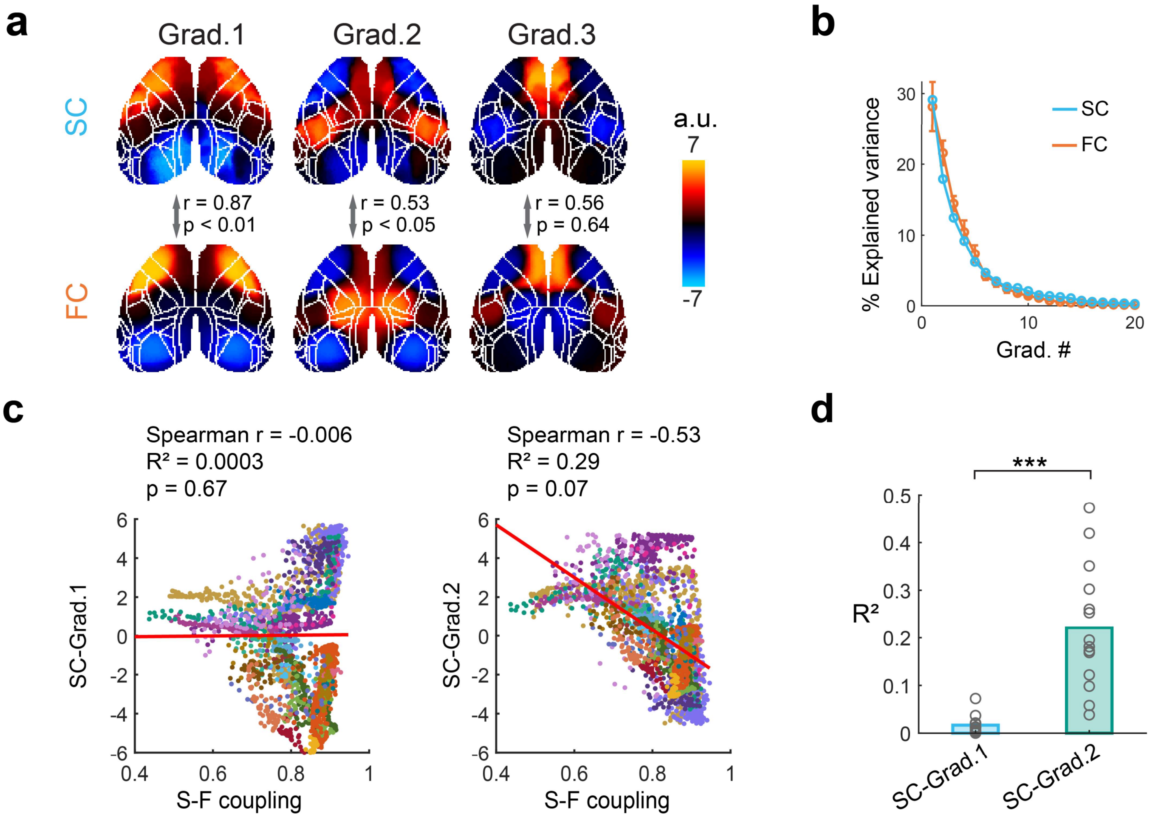


**Supplementary Fig. 5. Structure-Function Coupling Followed Gradient 2 Derived from Structural Connectivity**

**a)** Principal gradients derived from SC and FC. FC are reproduced from **Fig. 5a** for comparison. We calculated SC gradients in the same way as the FC gradients[7]. The Pearson correlations between corresponding pairs are shown. Significance of spatial correlations were assessed using the Moran permutation test. **b)** Variance explained by the top 20 principal functional and structural gradients. Data of functional gradients are presented as mean ± s.d.. **c)** Similar to **Fig. 5c**. Scatter plots comparing the S-F coupling against the scores of SC Gradients 1 and 2. Each dot represents a pixel on the cortex. Spatial correlations were assessed using the Moran permutation test. **d)** Similar to **Fig. 5d**. Comparison of correlation values between local S-F coupling and SC Gradients 1 and 2 for each individual animal ($n = 14$). S-F coupling is significantly more correlated with SC Gradient 2 than with SC Gradient 1 ($p<0.001$, paired t-test).

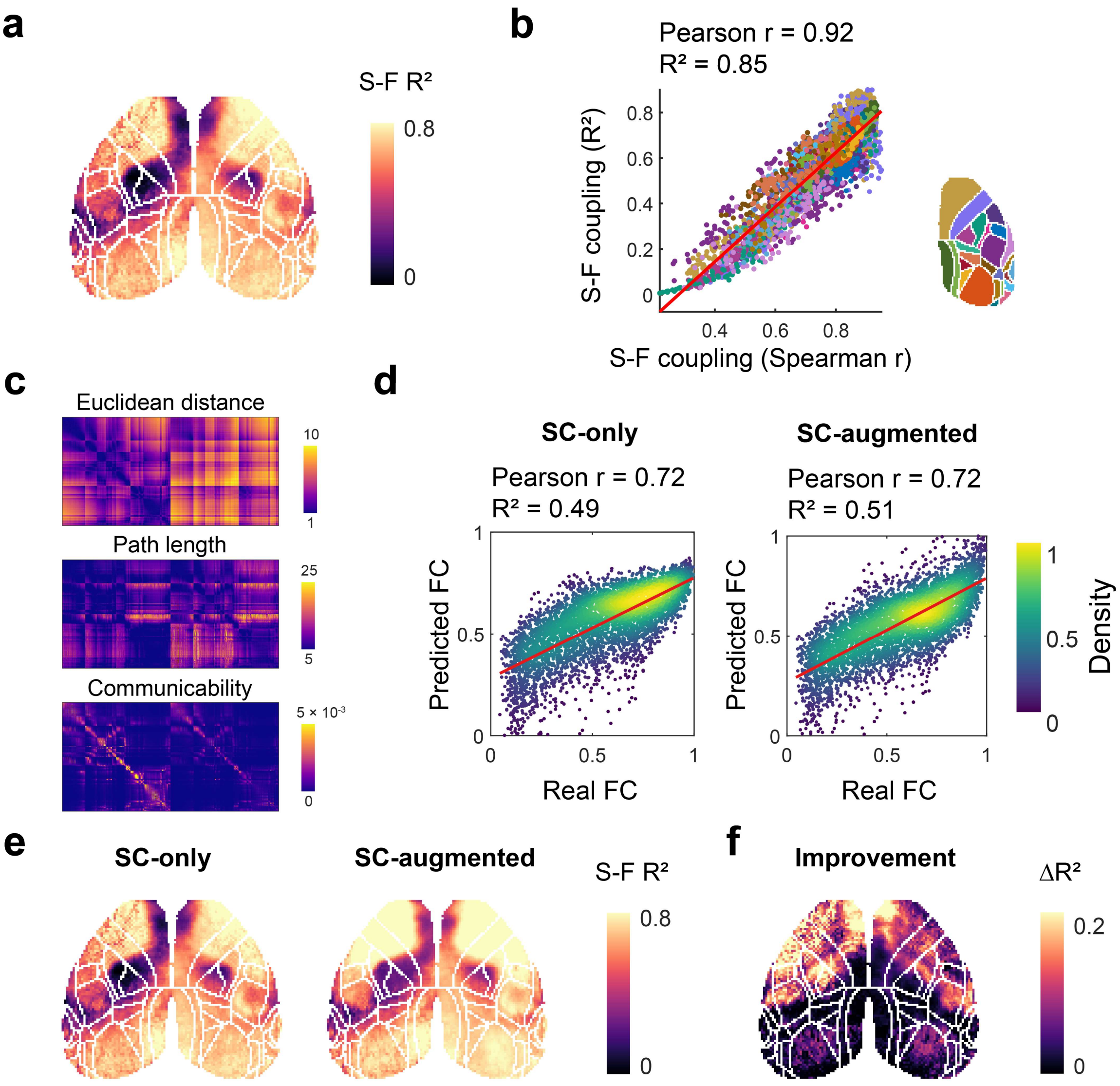


**Supplementary Fig. 6. Multiple Linear Regression in Evaluating S-F coupling**

**a)** Cortical map of local S-F coupling calculated using multiple linear regression as described previously[4,10] . The correspondence between SC and FC is reported as coefficient of determination of the model (S-F $R^2$). **b)** Comparing the local S-F $R^2$ and the S-F coupling estimated in our primary analysis. Each dot represents a cortical pixel and is color-coded by cortical area. **c)** Three additional structural features are calculated following methodologies from previous studies[4] . **d)** Comparison of SC-only and SC-augmented models' performance in predicting global FC. The plots show the correlation between observed FC and predicted FC when the predictive model used either SC alone (SC-only) or SC together with the additional structural features from (**c**) as predictors (SC-augmented). **e)** Map of local S-F $R^2$, calculated as in (**b**), either using only

SC alone or additional structural features from (**d**) as predictors. **f)** A map showing the difference in S-F $R^2$ values between two maps in **(e)**. This visualization highlights the improvement in explaining local FC variance gained by including the additional structural features in the model.
